# Using Computer-vision and Machine Learning to Automate Facial Coding of Positive and Negative Affect Intensity

**DOI:** 10.1101/458380

**Authors:** Nathaniel Haines, Matthew W. Southward, Jennifer S. Cheavens, Theodore Beauchaine, Woo-Young Ahn

## Abstract

Facial expressions are fundamental to interpersonal communication, including social interaction, and allow people of different ages, cultures, and languages to quickly and reliably convey emotional information. Historically, facial expression research has followed from discrete emotion theories, which posit a limited number of distinct affective states that are represented with specific patterns of facial action. Much less work has focused on dimensional features of emotion, particularly positive and negative affect intensity. This is likely, in part, because achieving inter-rater reliability for facial action and affect intensity ratings is painstaking and labor-intensive. We use computer-vision and machine learning (CVML) to identify patterns of facial actions in 4,648 video recordings of 125 human participants, which show strong correspondences to positive and negative affect intensity ratings obtained from highly trained coders. Our results show that CVML can both (1) determine the importance of different facial actions that human coders use to derive positive and negative affective ratings, and (2) efficiently automate positive and negative affect intensity coding on large facial expression databases. Further, we show that CVML can be applied to individual human judges to infer which facial actions they use to generate perceptual emotion ratings from facial expressions.

## Using Computer-vision and Machine Learning to Automate Facial Coding of Positive and Negative Affect Intensity

The ability to effectively communicate emotion is essential for adaptive human function. Of all the ways that we communicate emotion, facial expressions are among the most flexible, allowing us to rapidly convey information to people of different ages, cultures, and languages. Further, facial expressions signal complex action tendencies including threat and cooperative intent (1-3). Unsurprisingly, the ability to produce and recognize facial expressions of emotion is of interest to researchers throughout the social and behavioral sciences.

Facial expressions can be interpreted using either message- or sign-based approaches (4). Message-based approaches describe the meaning conveyed by a facial expression (e.g., happiness), whereas sign-based approaches describe observable facial actions that embody/comprise messages (e.g., cheek raising may indicate happiness). Although message-based approaches are used effectively by psychologists to measure facial expression messages (e.g., happiness), they do not describe facial behavior comprehensively. Instead, they rely on expert judgments of holistic facial expressions—provided by highly trained coders—rather than on facial movements themselves. This renders message-based approaches susceptible to individual differences (unreliability) among human coders, which can impede valid comparisons of results across studies and research sites—even when the same construct is measured.

In comparison, multiple comprehensive, standardized sign-based protocols have been developed and used to answer a variety of research questions (4). Among these protocols, the Facial Action Coding System (FACS; 5) may be the most widely used. FACS comprises approximately 33 anatomically-based facial actions (termed action units [AUs]), which interact to generate different facial expressions.

Originally developed from a basic emotion theory perspective, the relation between FACS-based AUs and discrete emotions is an active research topic (6). Distinct patterns of AUs reliably map onto each basic emotion category (happiness, sadness, anger, fear, surprise, and disgust), and the existence of distinct patterns of AUs that people use to label different emotional expressions is often used as evidence to support discrete theories of emotion (see 7). For example, oblique lip-corner contraction (AU12), together with cheek raising (AU6) reliably signals enjoyment (8), while brow furrowing (AU4) tends to signal negative emotions like anger and sadness (e.g., 9). Recently, research on how people perceive discrete emotions from AUs has revealed up to 21 discrete categories composed of compound basic emotions (e.g., happily-surprised; 10). Together, these studies suggest that people use the presence of distinct AUs to evaluate emotional content from facial expressions (11), a hypothesis supported by neuroimaging studies showing that differential patterns of BOLD responding in the posterior superior temporal sulcus discriminate between AUs (12).

Despite the clear links between AUs and discrete emotion perception, little is known about how AUs map onto dimensional features of emotion (7), especially positive and negative affect (i.e., valence). This is a potentially important oversight given the centrality of valance to dimensional theories of emotion (e.g., 13-15), of which valence is the most consistently replicated dimension (16). Early work using facial electromyography (EMG) showed that zygomatic (AU12) and corrugator (AU4) activity may indicate more positive and more negative subjective intensity, respectively (e.g., 9). However, later studies found that interactions between multiple AUs better describe valence intensity (e.g., 17), and in follow-up work, researchers have proposed that the face may represent positive and negative affect simultaneously with independent sets of AUs (e.g., 18). In one of the few studies directly linking AUs to perceived valence intensity, Messinger et al. (19) found that cheek raising (AU6) was common to perceptual judgments of both intense positive and negative affect, which challenges the idea that a single AU can adequately capture the entire range of valence intensity. Altogether, current evidence suggests that zygomatic (AU12) and corrugator (AU4) activity indicate perceived positive and negative affect, but the extent to which these and other discrete facial actions map onto the entire range of perceived positive or negative affect intensity is unclear.

Comprehensive follow-up investigations have been difficult to pursue, in part, because measuring AUs is labor- and time-intensive and requires highly skilled annotators. Indeed, FACS training requires an average of 50-100 hours, and minutes of video can take expert coders multiple hours to rate reliably (20). These characteristics limit sample sizes, reduce feasibility of replication efforts, and discourage researchers from coding facial expressions. Instead, researchers tend to rely on measures of emotional responding that are not observable in social interactions (e.g., heart rate variability). Recently, automated computer-vision and machine learning (CVML) based approaches have emerged that make it possible to scale AU annotation to larger numbers of participants (e.g., 21-23) thus making follow-up studies more feasible. In fact, inter-disciplinary applications of CVML have allowed researchers to automatically identify pain severity (e.g., 24), depressive states (e.g., 25), and discrete emotions from facial expressions (e.g., 26).

Work using CVML to detect valence intensity from facial expressions is ongoing (see 27). In fact, there are annual competitions held to develop CVML models that best characterize dimensional features of emotions such as valence and arousal (e.g., 28). Currently, basic emotions can be coded automatically with accuracy comparable to human coders, but valence intensity models show lower concurrent validity. For example, state-of-the-art CVML models show correlations between human- and computer-coded valence ranging from *r* = .60-.71 (29,30). While impressive, these CVML models are often constructed to detect valence directly from frame-by-frame video input without intermediately capturing AUs, so it is unclear if successful valence detection depends on prior detection of specific AUs. Furthermore, these results have been collected from relatively small samples or on continuously collected valence ratings (human ratings collected in real-time using dials or joysticks), and it is unclear if these models generalize to other research settings where participants’ emotional expressions to evocative stimuli are coded within discrete, trial-by-trial time intervals (e.g., 31). Indeed, contemporary work using CVML has shifted from evaluating facial expressions in controlled laboratory settings toward accurately capturing continuous facial expressions of emotion “in the wild”, which is a much more difficult task (e.g., 29,32). However, to be useful for the majority of applications within the cognitive, decision, and other social and behavioral sciences—where controlled laboratory settings are the norm—it is only necessary that CVML models are optimized for performances within laboratory settings as opposed to in the real-world. Lastly, most valence-detecting CVML models assume a unidimensional valence continuum as opposed to separable continua for positive and negative affect—to our knowledge, there are few opensource datasets used in CVML research that characterize valence as multi-dimensional (see 33), and very little work has been done with CVML to separate positive and negative affect (cf. 34). Notably, positive and negative affect can vary independently and have different predictive values (10,15,35), suggesting that CVML models designed to account for each dimension separately may be most beneficial for behavioral science applications.

Using a well-validated method of emotion induction and both computer-vision measurement of discrete facial actions and continuous measures of positive and negative affect intensity, we (1) identified specific correspondences between perceived emotion intensity and discrete facial AUs, and (2) developed a reliable, valid, and efficient method of automatically measuring the separable dimensions of positive and negative affect intensity. Importantly, data used to train and validate our CVML models were collected from a commonly-used psychological task and contained 4,648 video-recorded, evoked facial expressions from 125 human subjects across multiple task instructions. Our findings shed light on the mechanisms of valence recognition from facial expressions and point the way to novel research applications of large-scale emotional facial expression coding.

## Method

### Participants

Video recordings and human coder data were collected as part of a larger study (31). The current study included 125 participants (84 females), ages 18-35 years. All participants gave informed consent prior to the study, and the study protocol (#2011B0071) was approved by The Ohio State Behavioral and Social Sciences Institutional Review Board. Self-reported ethnicities of participants were as follows: Caucasian (*n*=96), East Asian (*n*=14), African-American (*n*=5), Latino (*n*=3), South Asian (*n*=3), and unspecified (*n*=4).

### Measures

#### Emotion-evoking task

We used an emotion-evoking task, depicted in Figure 1, that has been used in several previous studies to elicit facial expressions of emotion across multiple task instructions (31,36). Participants viewed 42 positive and negative images selected from the International Affective Picture System (IAPS) to balance valence and arousal. Selections were based on previously reported college-student norms (37). Images were presented in 6 blocks of 7 trials each, whereby each block consisted of all positive or all negative images. For each block, participants were asked to either *enhance*, *react normally*, or *suppress* their naturally evoked emotional expressions to the images. These instructions effectively increased variability in facial expressions within participants, and they reflect common social situations where emotions must be enhanced or suppressed to achieve desired outcomes. Given known individual differences in suppression and enhancement of facial expressions (31,36), we expected that these task instructions would allow us to create a more generalizable CVML model than with no instructions at all. Block order was randomized across participants. Instructions were given so that each valence was paired once with each condition. All images were presented for 10 s, with 4 s between each image presentation. Participants’ reactions to each image were video-recorded with a 1080p computer webcam (Logitech HD C270). Due to experimenter error, 1 participant’s videos were not recorded correctly, and 7 participants were shown only 41 recordings, resulting in 6,293 usable recordings. Among these, 3 were corrupted and could not be viewed. Thus, 6,290 10-s recordings were potentially available.

**Figure 1.**
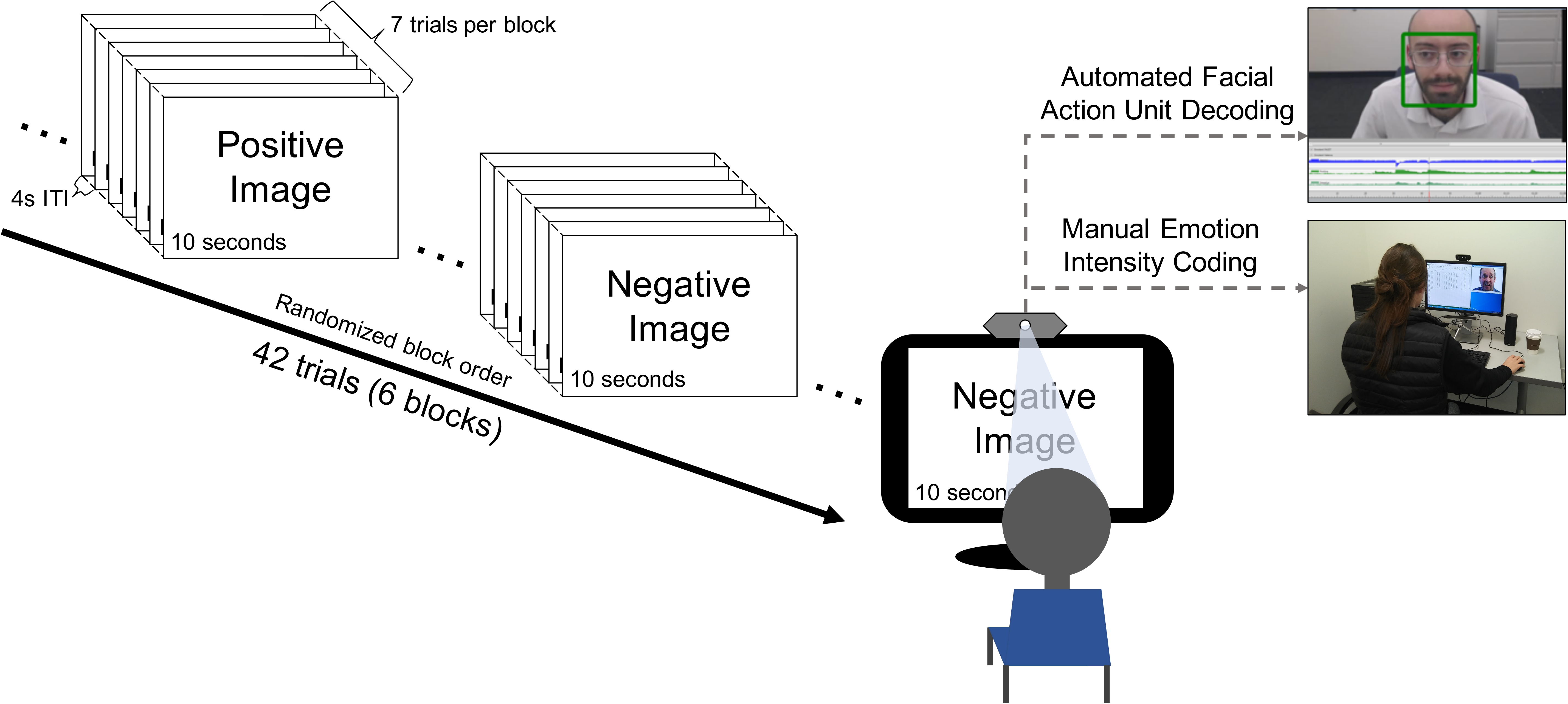
Emotion-evoking task. Participants (*N*=125) viewed a total of 42 images each, divided into 6 blocks of 7 trials. Images were presented for 10 s, with a 4 s inter-trial-interval. Each block of images consisted of either positive or negative image content. In each of the 3 blocks containing positive and negative image content, participants were asked to either *enhance*, *react normally*, or *suppress* their emotional expressions, so that each valence type (i.e., positive or negative) was paired once with each task instruction (enhance, react normally, suppress). All images were selected from the International Affective Picture System (37). Participants’ reactions to the images were video recorded and their facial expressions were subsequently rated for positive and negative emotion intensity by a team of trained coders. The same recordings were then analyzed by FACET, a computer vision tool which automatically identifies facial Action Units (AUs).

#### Manual coding procedure

A team of three trained human coders, unaware of participants’ task instructions, independently viewed and rated each 10-s recording for both positive and negative emotion intensity. Presentation of recordings was randomized for each coder. Ratings were collected on a 7-point Likert scale ranging from 1 (*no emotion*) to 7 (*extreme emotion*). Coders completed an initial training phase during which they rated recordings of pre-selected non-study cases and discussed specific facial features that influenced their decisions (see the Supplementary Text for the coding guide). The goal of this training was to ensure that all coders could reliably agree on emotion intensity ratings. In addition, coders participated in once-monthly meetings throughout the coding process to ensure reliability and reduce drift. Agreement between coders across all usable recordings (6,290 recordings) was high, with intraclass correlation coefficients (ICCs(3); 38) of .88 and .94 for positive and negative ratings, respectively. The ICC(3) measure reported above indicates absolute agreement of the average human-coder rating within each condition (*enhance*, *react normally*, *suppress*) for each of the 125 participants. To prepare data for CVML analysis, we performed an additional quality check to screen out videos in which participants’ faces were off-camera or covered. Any recording in which a participant’s face was covered, obscured, or off-camera for 1 s or more was removed from analysis. If 50% or more of a participant’s recordings were excluded, we excluded all of his/her recordings. This resulted in a total of 4,648 usable recordings across 125 participants. With over 4,000 individually-coded recordings, our sample size is in the typical range for machine learning applications (39).

#### Automated coding procedure

We then analyzed each of the 4,648 recordings with FACET (23). FACET is a computer-vision tool that automatically detects 20 FACS-based AUs (see Supplementary Table 1 for descriptions and depictions of FACET-detected AUs). FACET outputs values for each AU indicating the algorithm’s confidence in the AU being present. Confidence values are output at a rate of 30 Hz, resulting in a time-series of confidence values for each AU being present with each frame of a video-recording. Each point in the time-series is a continuous number ranging from about −16 to 16, whereby more positive and more negative numbers indicate increased and decreased probability of the presence of a given AU, respectively. We refer to this sequence of numbers as an AU evidence time-series.

Each AU evidence time-series was converted to a point estimate by taking the area under the curve (AUC) of the given time-series and dividing the AUC by the total length of time that a face was detected throughout the clip. This creates a normalized measure that does not render biased weights to clips of varying quality (e.g., clips in which participants’ faces are occasionally not detected). Point-estimates computed this way represent the expected probability that a participant expressed a given AU across time. We used the AU evidence time-series point estimates as predictor (independent) variables to train a machine learning model to predict human valence intensity ratings. It took FACET less than 3 days to extract AU evidence time-series data from all recordings (running on a standard 8-core desktop computer). Note that we did not use a baseline correction for each subject, which would require human annotation of a neutral facial expression segment for each participant. Therefore, the models reported here may be applied to novel facial recordings with no human judgment.

#### Machine Learning Procedure

Figure 2 depicts the machine learning procedure. We trained a random forest (RF) model to predict human-coded valence ratings from the AU evidence time-series point estimates described above (see Supplementary Text for details on training). RFs are constructed by generating multiple decision trees and averaging predictions of all trees together. We chose the RF model because (1) it can automatically capture interactions between independent variables, and we know that humans use multiple AUs simultaneously when evaluating facial expressions; (2) the importance of each independent variable can be extracted from the RF to make inferences regarding which AUs human coders attended to while rating valence intensity (analogous to interpreting *beta* weights from a multiple regression; 39); and (3) RFs have previously shown robust representations of the mapping from facial features (e.g., AUs) to discrete emotions and valence intensity (40,41). Given high agreement among coders and a large literature showing that aggregating continuous ratings from multiple, independent coders leads to reliable estimates despite item-level noise (i.e., ratings for each recording; see 42), we used the average of all coders’ ratings for each recording as the outcome (dependent) variable to train the RF.

**Figure 2.**
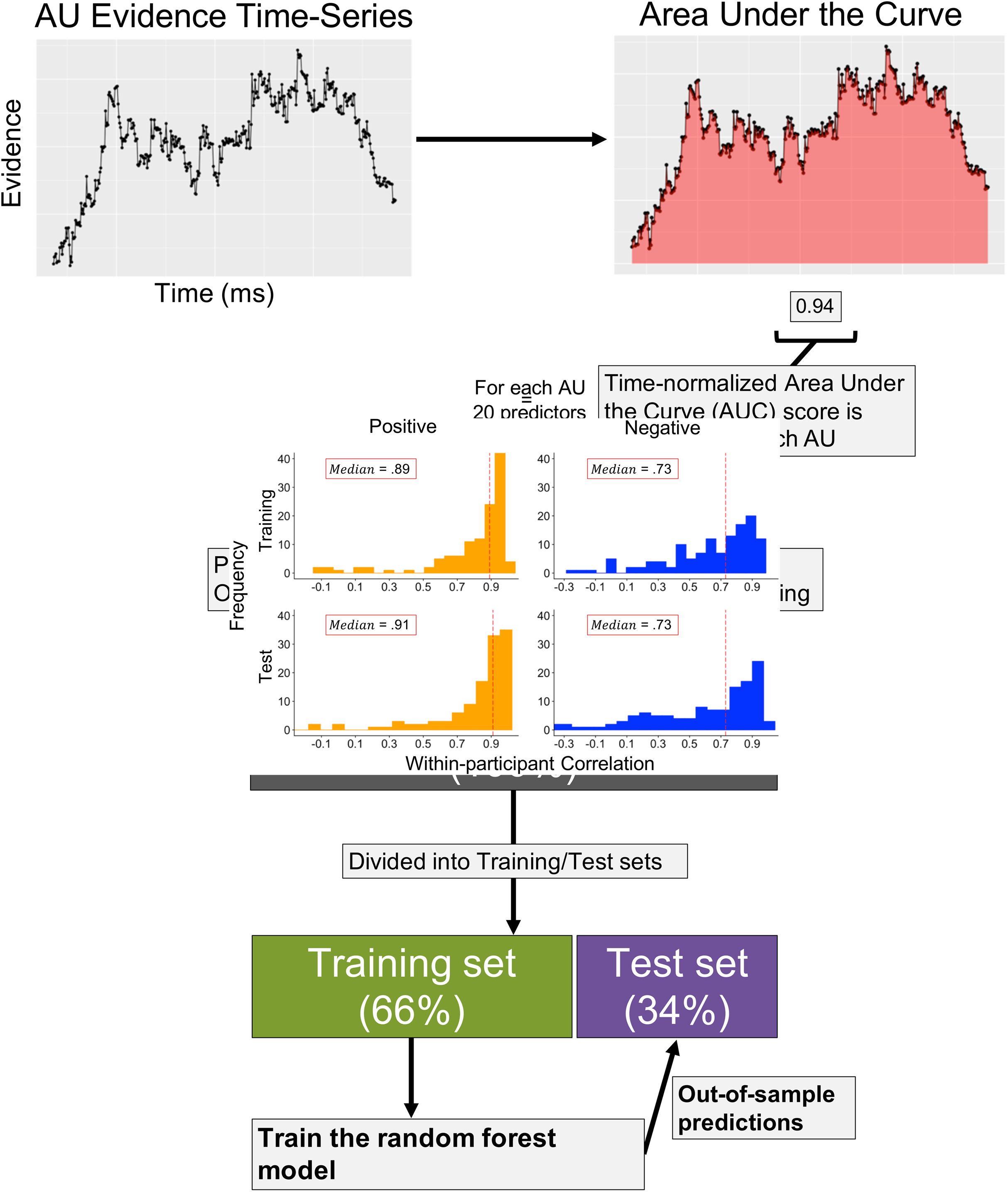
Machine learning procedure. The goal of our first analysis was to determine whether or not CVML could perform similarly to humans in rating facial expressions of emotion. For each AU evidence time-series, we computed the normalized (i.e., divided by the total time that FACET detected a face) Area Under the Curve (AUC), which captures the probability that a given AU is present over time. All AUC values (20 total) were entered as predictors into the random forest (RF) model to predict the average coder rating for each recording. To test how similar the model ratings were to human ratings, we separated the data into training (3,060 recordings) and test (1,588 recordings) sets. We fit the RF to the training set and made predictions on the unseen test set. Model performance was assessed by comparing the Pearson and intraclass correlations between computer- and human-generated ratings in the test sets.

The RF model contains 2 tuning parameters, namely: 1) *ntrees*–the number of decision trees used in the forest, and 2) *mtry*–the number of predictors to sample from at each decision node (i.e., “split”) in a tree. A grid search over *ntrees ∈* {100, 200, 300, …, 1000} showed that out-of-bag prediction accuracy converged by 500 trees for both positive and negative datasets (not reported). A grid search over *mtry ∈* {1, 2, 3, …, 20} revealed negligible differences in out-of-bag prediction accuracy for values ranging from 5 to 20. Because RFs do not over-fit the data with an increasing number of trees (39), we set *ntrees* = 500 for models presented in all reported analyses to ensure convergence. Because initial grid searches over *mtry* failed to improve the model, we set *mtry* heuristically (39) as *mtry* = *p*/3, where *p* represents the number of predictors (i.e., 1 for each AU) in an *n* × *p* matrix (*n* = number of cases) used to train the model. We fit the RF model using the *easyml* R package (43), which provides a wrapper function for the *randomForest* R package (44). All R codes used for model fitting along with the trained RF models will be made available on our GitHub repository upon publication (https://github.com/CCS-Lab).

#### Correspondence between human coders and model predictions

Model performance refers to how similar the model- and human-generated valence intensity rating are. To assess model performance, we split the 4,648 recordings into training (*n*=3,060; 65.8%) and test (*n*=1,588; 34.2%) sets, trained the model on the training set (see the Supplementary Text for details), and then made predictions on the unseen test set to assess how well the RF predicted valence intensity ratings on new data. The data were split randomly with respect to participants so that the training and test data contained 66% and 34% of each participant’s recordings, respectively. This separation ensured that training was conducted with all participants, thus creating a more generalizable final model. We fit a separate RF model to positive and negative human ratings. To see if the way we split the training and test data influenced our results, we made 1,000 different training/test-set splits and assessed model performance across all splits (45,46). We used Pearson correlations and ICC coefficients to check model performance on training- and test-sets. Pearson correlations measure the amount of variance in human ratings captured by the model, whereas ICCs measure absolute agreement between human- and model-predicted ratings at the item level (i.e., per recording). Therefore, high correlations and ICCs indicate the model is capturing a large amount of variance in human coder ratings and generating ratings using a similar scale as human coders, respectively. We used McGraw and Wong’s ICC(1), as opposed to other ICC methods (38), because we were interested in absolute agreement across all clips, regardless of condition/participant. One-way models were used to compute ICCs in all cases. In general, ICCs between .81 and 1.00 are considered “almost perfect” (i.e., excellent) and ICCs between .61 and .80 are considered “substantial” (i.e., good; 47). We also checked model performance using a different folding scheme for separating training and test sets which ensured that participants’ recordings were not shared across splits. This analysis revealed negligible differences in prediction accuracy for positive ratings and a decrease in accuracy for negative ratings, which suggests that more training data may be necessary to capture negative as opposed to positive affect intensity (see Supplementary Text).

#### Importance of AUs for positive and negative affect

To identify the specific AUs that human coders were influenced most by when making affective ratings, we fit the RF model to the entire dataset (all 4,648 recordings) without splitting into training and test sets. We used this method to identify independent variables that were robust across all samples (45,46). After fitting the RF models, the importance of each independent variable was estimated using *increase in node purity*, a measure of change in residual squared error (increases in prediction accuracy) attributable to the independent variable across all decision trees (39). Importance scores for each independent variable extracted from the RF then represent the magnitude—but not direction of— the effect a given AU has on human coders’ valence ratings.

To identify potential individual differences in emotion recognition between human coders, we also fit the RF to each coder’s ratings independently. We used randomization tests to determine the minimum number of ratings necessary to accurately infer which AUs the coders attended to while generating emotion ratings. For each of the 3 coders, we performed the following steps: (1) randomly sample *n* recordings rated by coder *i*, (2) fit the RF model to the subset of *n* recordings/ratings according to the model fitting procedures outlined above, (3) compute the ICC(2) of the extracted RF feature importances (i.e., *increase in node purity*) between the permuted model and the model fit to all recordings/ratings from coder *i*, and (4) iterate steps 1-3 twenty times for each value of *n* (note that different subsets of *n* recordings/ratings were selected for each of these twenty iterations). We varied *n* E {30, 40, 50, 60, 70, 80, 90, 100, 200, 300, 400, 500, 600, 700, 800, 900, 1000, 1200, 1400, 1600, 1800, 2000, 2500, 3000}.

## Results

### Model performance across participants

Table 1 shows correlations between the model-predicted and the average of the human coders’ ratings per recording across both training and test sets. Overall, the RF showed good to excellent performance across both training and test sets for positive and negative ratings. Notably, these results were supported by both the Pearson correlations and the ICCs, suggesting that the RF produced ratings that not only captured variance in, but also showed high agreement with, human ratings. Sensitivity analyses (see Figure 3) indicated that model performance was robust across different training and test splits of the data. These results suggest that variance in human-coded valence intensity can be captured by the presence of discrete AUs.

**Table 1.** Correlations between human- and computer-generated valence ratings.

| Data Set | Correlation [95% CI] |  |  |  |
| --- | --- | --- | --- | --- |
| | $r$ | | ICC(1) | |
|  | (+) | (-) | (+) | (-) |
| Training | .89 [.88, .90] | .77 [.75, .78] | .88 [.87, .89] | .71 [.69, .72] |
| Test | .88 [.87, .89] | .74 [.72, .77] | .87 [.86, .88] | .68 [.65, .71] |

*Notes.* (+) = positive valence ratings; (-) = negative valence ratings; $r$ = Pearson's correlation; ICC = Intraclass correlation coefficient. Training and test sets contained 3,060 and 1,588 recordings, respectively.

**Figure 3.**
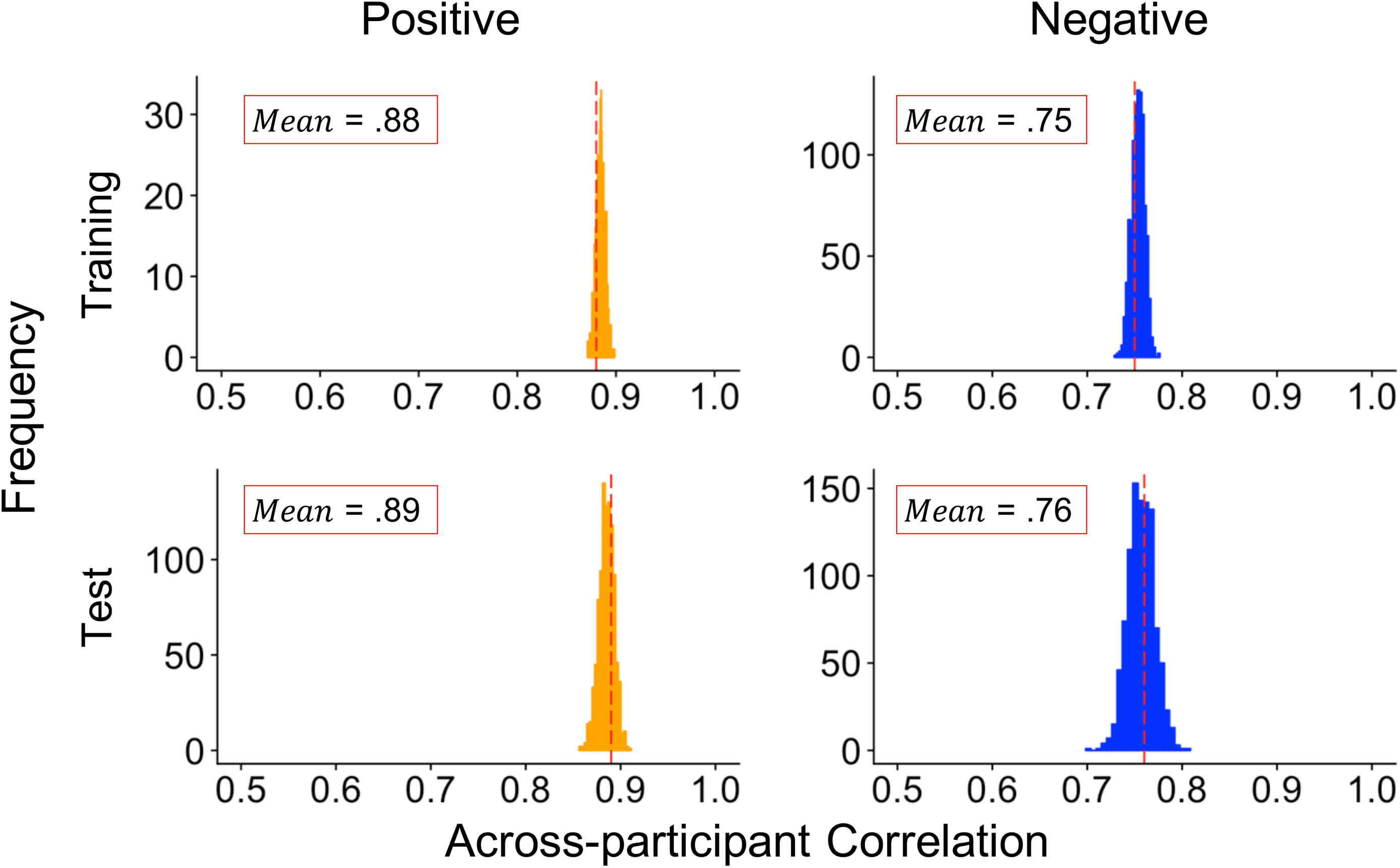
Sensitivity of model performance to different training/test splits. Results of sensitivity analyses across different splits of the training and test sets. We created 1,000 different splits of the training and test sets, fit the RF to each training set, and then made predictions on each respective test set. We stored the Pearson correlations between human- and model-generated ratings for each iteration. Distributions therefore represent uncertainty in prediction accuracy. Means of the distributions (superimposed on respective graphs) are represented by dashed red lines.

### Model performance within participants

We also checked model performance for each of the 125 participants by computing correlations between human- and model-generated ratings for each participant separately (Figure 4). Although the RF model performed well for many participants in the positive (median *r* = .91, ICC(1) = .80) and negative (median *r* = .73, ICC(1) =.51) affect test sets, 5 participants within the positive and 7 participants within the negative affect test-set yielded negative correlations between human- and computer-generated emotion ratings (Figure 4). Further analyses of within-participant model performance revealed significant positive associations between within-subject variance in model-predicted ratings and within-participant prediction accuracy (all *r*s :2 .54, *p*s < .001; see Supplementary Figure 1A). We found the same relation between human-assigned ratings and within-participant variance (see Supplementary Figure 1B). This suggests that the RF model was more accurate in predicting human-rated emotion if participants expressed a wider range of emotional intensity.

**Figure 4.**
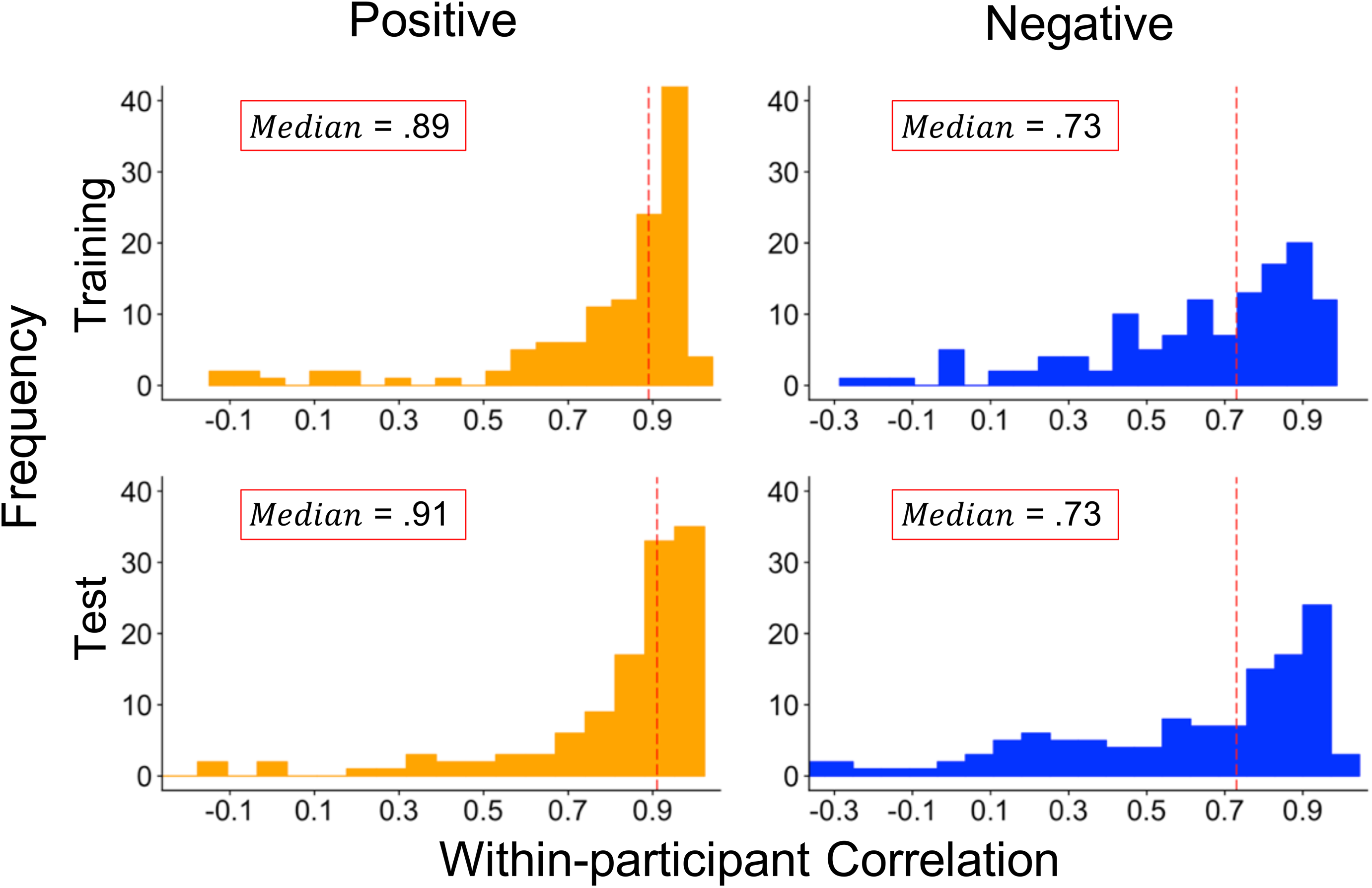
Model performance within participants. Distributions of within-participant Pearson correlations for positive and negative ratings in the training (all 125 participants) and test (122 participants; correlations could not be computed for 3 participants who had 0 variance in human ratings) sets. Red dashed lines represent median within-participant Pearson correlations for each distribution. Intraclass correlations for corresponding figures are reported in text.

### Importance of AUs across task instructions

To identify which facial expressions human coders may have used to generate positive and negative emotion ratings, we examined the importance of all AUs in predicting human emotion ratings (Figure 5). Note that importance values for the RF do not indicate directional effects, but instead reflect relative importance of a given AU in predicting human-coded positive/negative affect intensity. The RF identified AUs 12 (*lip corner pull*), 6 (*cheek raiser*), and 25 (*lips part*) as three of the four most important AUs for predicting positive emotion. Together, AUs 12 and 6 accounted for 50% of the total importance (analogous to proportion of variance accounted for) of all AUs for positive ratings. In contrast to positive ratings, relative importance values for AUs of negative ratings were distributed more evenly across AUs, a trend which was also found when the RF was fit individually to each coder (see *Individual differences in emotion recognition* below). In fact, the 5 most important AUs for negative ratings (Figure 5) accounted for 50% of the total importance of all AUs, compared to only AU12 and AU6 accounting for the same proportion of importance in positive ratings. Notably, the importance of AUs for positive and negative emotion ratings were largely independent. In fact, when the ICC(3) is computed by treating positive and negative importance weights for each AU as averaged ratings from two “coders”, the ICC(3) is negative and non-significant (ICC(3) = –.11, *p* = .58), which would only be expected if different facial expressions were important for the coders to rate positive versus negative valence.

**Figure 5.**
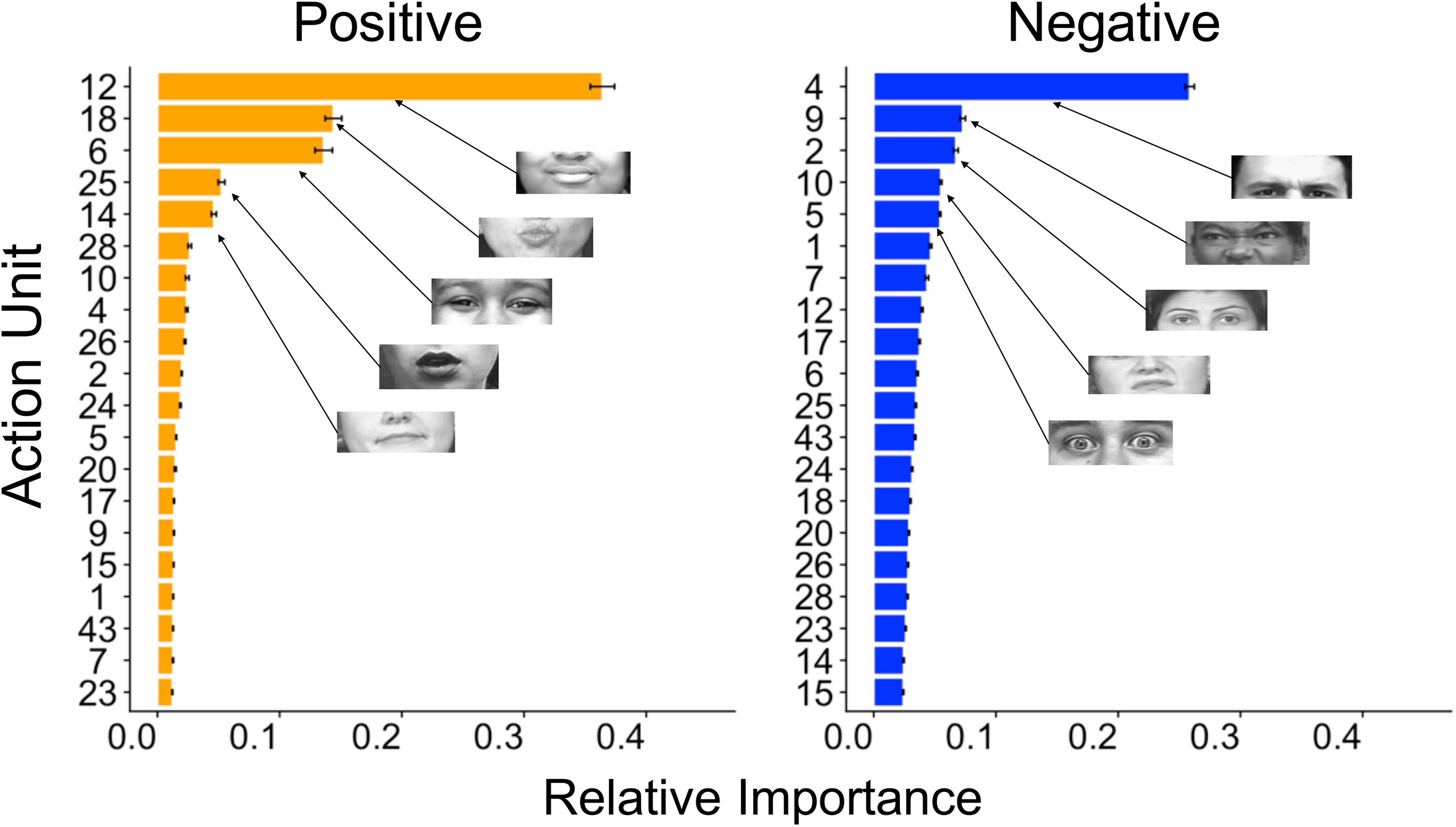
Relative importance of each AU for positive and negative ratings. Relative importance of each AU for positive and negative human-coder ratings. Relative importance (normalized *increase in node purity* from the RF model) is a measure of change in residual squared error, (increase in prediction accuracy) attributable to each AU being included in the model, and it can be interpreted as how important a given AU is with respect to all other AUs. Note that relative importance is not directional, but does capture interactions among AUs. Visual depictions of the 5 most important AUs for predicting positive and negative ratings are shown on the graphs. Error bars indicate 95% confidence intervals (CIs) estimated from the sensitivity analysis (i.e., 1,000 train/test splits).

### Sensitivity of AUs to task instructions

To determine if task instructions (*enhance*, *react normally*, *suppress*) affected model performance or our interpretation of which AUs map onto positive and negative affect, we fit the RF model to all recordings from each condition separately and then compared model performance and AU importance scores across conditions. Table 2 shows correlations between human- and computer-generated valence ratings within the different conditions. For positive ratings, correlations were consistently high (*r*s > .80) across all conditions. In contrast, for negative ratings, correlations were highest in the enhance condition, followed by the react normally and suppress conditions. Of note, all correlations between human- and computer-generated ratings were lower when data were separated by condition compared to when condition was ignored (cf., Table 2 to Table 1). This suggests the lower number of recordings included in the training samples may be partially responsible for lower model performance, but also that CVML performs best when trained on a wider range of emotional intensity.

**Table 2.** Correlations between human- and computer-generated ratings within conditions.

| Condition | Correlation [95% CI] |  |  |  | Number of recordings |  |
| --- | --- | --- | --- | --- | --- | --- |
|  | *r*<br>(+) | *r*<br>(-) | ICC(1)<br>(+) | ICC(1)<br>(-) | Training | Test |
| Enhance | .81 [.78, .84] | .64 [.59, .68] | .79 [.76, .82] | .61 [.55, .66] | 1,047 | 569 |
| Normal | .81 [.78, .84] | .55 [.49, .61] | .79 [.76, .82] | .49 [.42, .55] | 880 | 516 |
| Suppress | .85 [.83, .87] | .44 [.38, .51] | .83 [.80, .85] | .35 [.28, .42] | 1,040 | 596 |

Notes. (+) = positive valence ratings; (–) = negative valence ratings; $r$ = Pearson’s correlation; ICC = Intraclass correlation coefficient. All results reported are on test sets.

Despite only moderate correlations for negative ratings in these conditions, relative importance values for AUs across conditions showed only small differences (Figure 6). In fact, ICCs between AU importance values across conditions were excellent for both positive and negative ratings (Figure 6). This suggests that the task instructions did not strongly influence which AUs were most important to coders.

**Figure 6.**
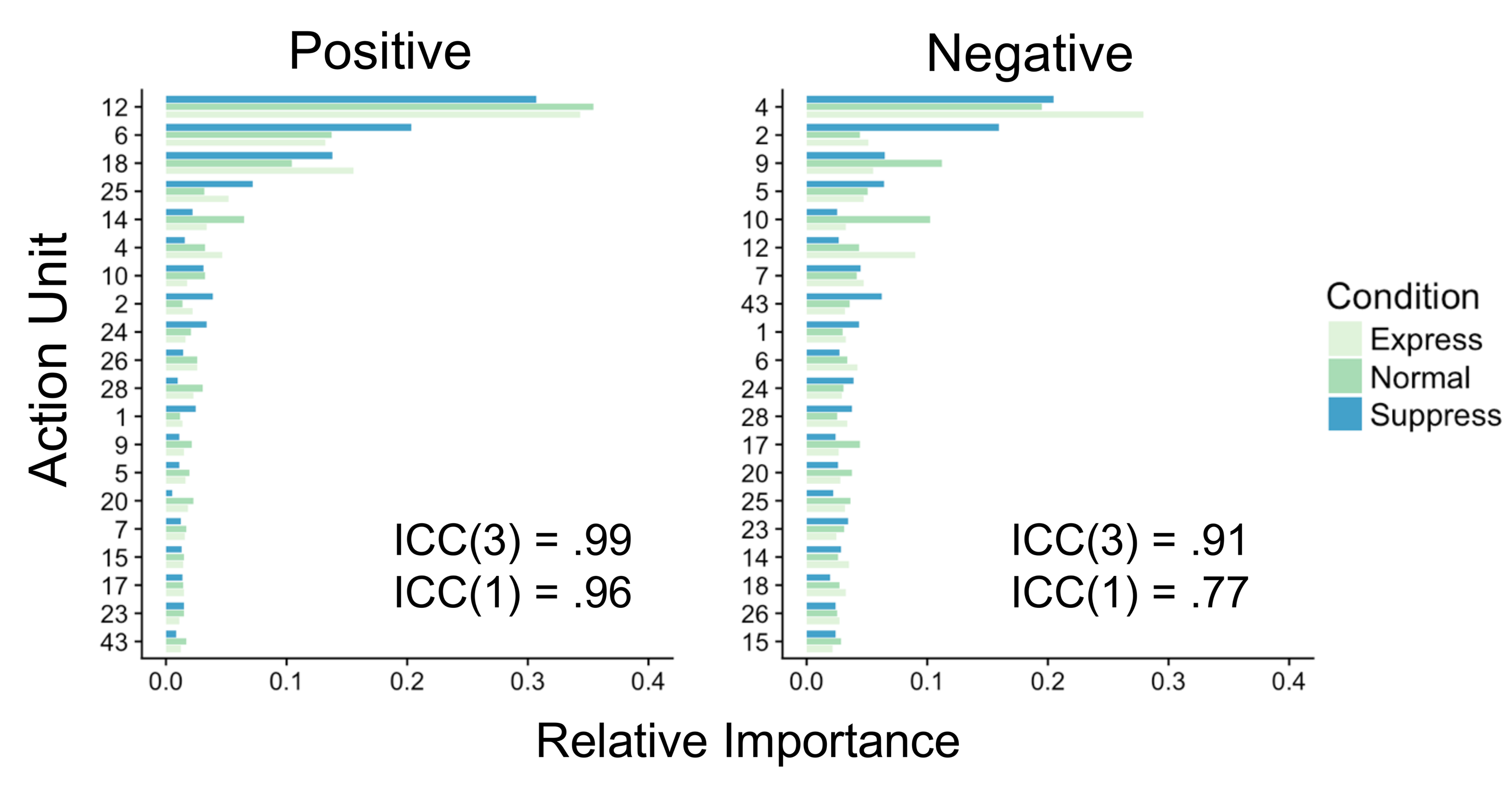
AU relative importance values across task instructions. Relative importance of each AU for positive valence and negative valence human-coder ratings within each of the three task instructions (*enhance*, *react normally*, *suppress*). Intraclass correlation coefficients–both treating importance values as average [ICC(3)] and single [ICC(1)] units–are superimposed. We show ICC(3) here because the AU importance scores could be interpreted as “averages” across all recordings.

### Individual differences in emotion recognition

All three coders showed similarly-ordered importance profiles, indicating that they attended to similar AUs while generating emotion ratings (Figure 7). Agreement between all three individual coders’ importance profiles supported this claim—ICC(3)s were high for both positive (ICC(3) = 0.95) and negative (ICC(3) = 0.93) importance profiles. Figure 8 shows the results of the randomization test. For positive ratings, ICC(2)s for all 3 coders reached 0.75 (regarded as “excellent” agreement; see 47) after just 60 recordings/ratings. For negative ratings, ICC(2)s for all 3 coders reached 0.75 after 200 recordings/ratings. Because the recordings in our task were 10 s long and coders rated positive/negative emotion intensity after each recording, the task used in the current study could be condensed to 200 recordings (~33 minutes) and still reveal individual differences in AU importances between coders with good accuracy. Future studies may be able to shorten the task even further by testing shorter video recordings (i.e., less than 10 s per recording).

**Figure 7.**
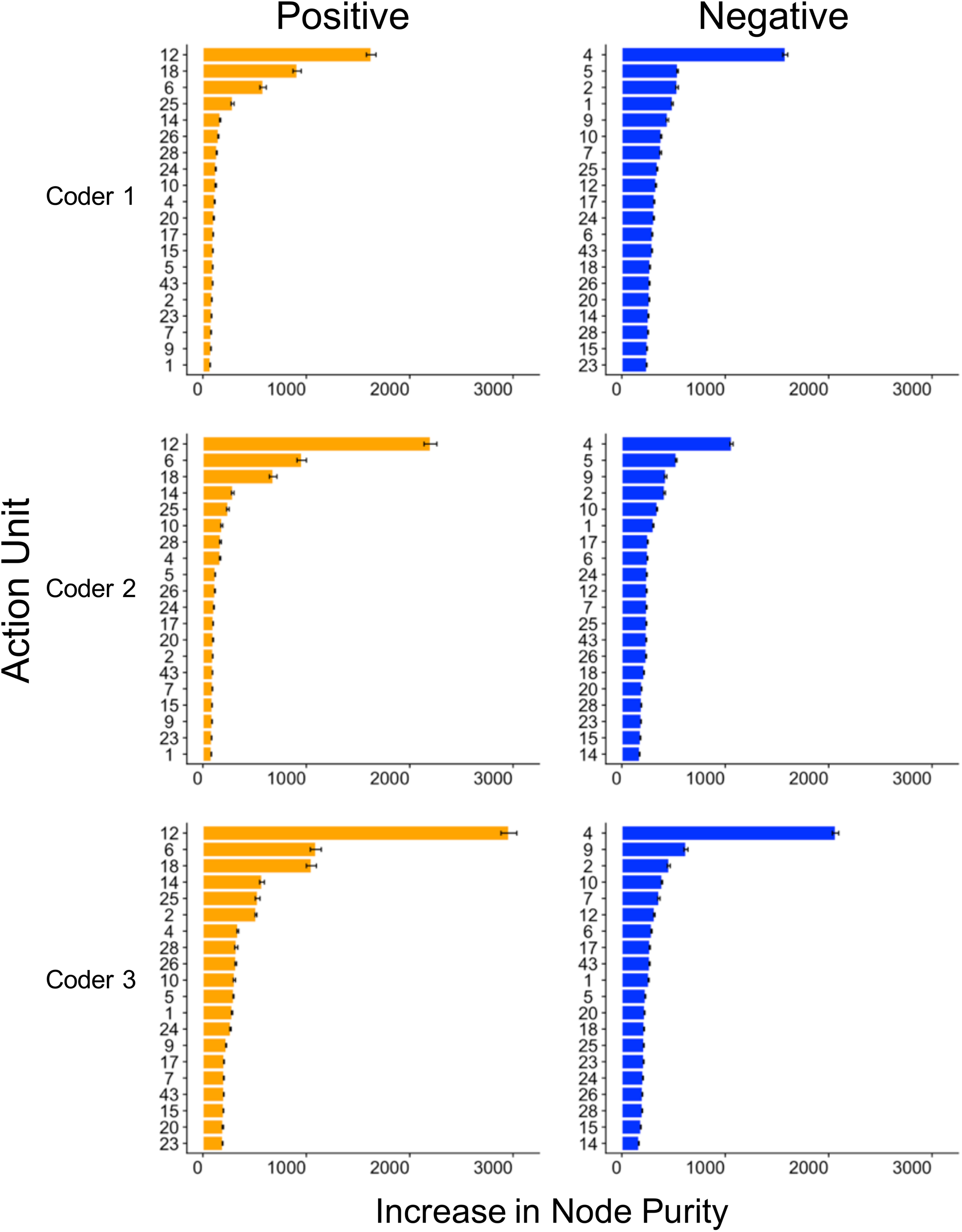
Individual differences in AU importances between coders. Feature importances (not normalized to show relative differences) extracted from the RF model fit separately to each coder. Coders all show similarly ordered importance profiles, suggesting that they attended to similar facial expressions while generating emotion ratings. Note that positive importance estimates are distributed across few predictors (i.e., AUs 6, 12, and 18), whereas negative importance estimates are more spread out throughout all predictors. Agreement between all three individual coders’ importance profiles was high, with ICC(3)s of .95 and .93 and ICC(1)s of .86 and .83 for positive and negative ratings, respectively.

**Figure 8.**
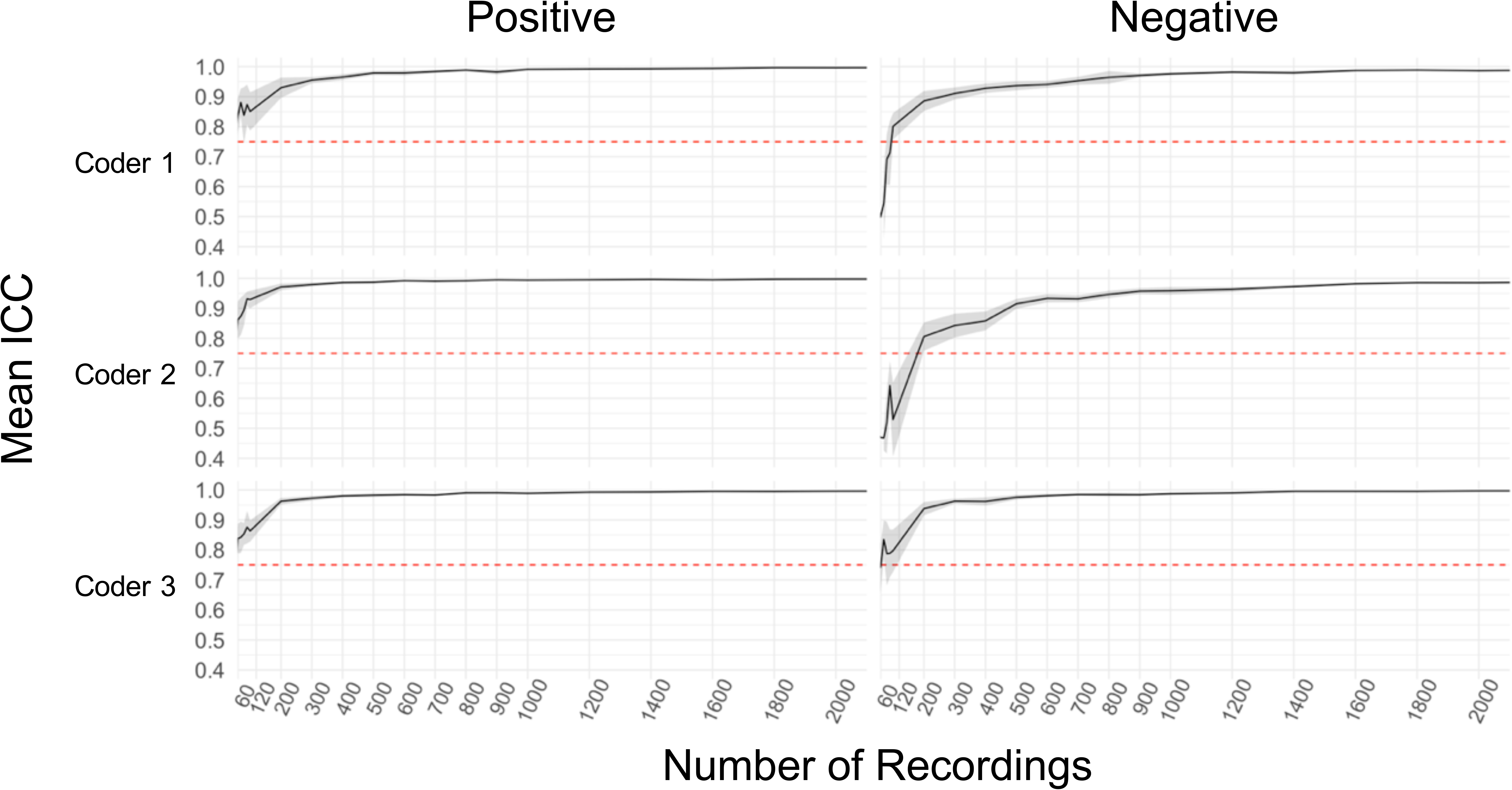
Number of recordings necessary to accurately estimate AU importance. Grid searches over the number of recordings/ratings necessary to achieve reliable estimates of AU importances for each valence-coder pair (coders appear in the same order as in Figure 7). Reliability is indexed by the ICC(2) between AU importance profiles (i.e. *increase in node purity*) extracted from the model fit to all the recordings that coders rated versus the model fit to subsets of recordings that they rated. Note that the ICC(2) assumes that importance estimates are “average” units (similar to ICC(3)s in Figures 6 & 7). The RF model was fit to each sample of size *n* along the *x*-axis, AU importance profiles were extracted from the model, and ICC(2)s were then calculated between the given sample and full-data AU importance profile scores. We iterated this procedure 20 times within each different sample size to estimate the variation in estimates across recordings. Shading reflects the 2 standard errors from the mean ICC within each sample across all 20 iterations. The red-dashed line indicates an ICC(2) of .75, which is considered “excellent”. For positive ratings, the ICC(2) reached .75 after only 60 recordings/ratings for each coder. For negative ratings, coders 1 and 3 reached an ICC(2) of .75 by 120 recordings/ratings, whereas coder 3 reached an ICC(2) of .75 by 200 recordings/ratings.

## Discussion

Our study offers strong evidence that people use discrete AUs to make judgments regarding positive and negative affect intensity from facial expressions, indicating that patterns of discrete AUs reliably represent dimensions of facial expressions of emotion (analogous to how specific patterns of AUs map to the basic emotions). Our CVML analysis identified AU12, AU6, and AU25 as especially important features for positive affect intensity ratings. Together, these AUs represent the core components of a genuine smile (48). Note that AU12 and AU6 interact to signify a *Duchenne smile*, which can indicate genuine happiness (8), and previous research demonstrates that accurate observer-coded enjoyment ratings rely on AU6 (49). Additionally, the five most important AUs we identified for negative affect intensity map on to those found in negative, discrete emotions such as anger (AUs 4 and 5), disgust (AU9), sadness (AU4), and fear (AUs 2, 4, and 5). Together, the AUs that we identified for positive and negative affect are consistent with prior studies suggesting that positive and negative facial expressions occupy separate dimensions (15,50). Notably, the AUs accounting for the majority of the variance in positive affect had no overlap with those for negative affect, evidenced by near-zero ICCs, indicating that our human coders used distinct patterns of facial expressions to evaluate positive versus negative intensity ratings. The existence of distinct patterns of AUs which represent positive and negative affect intensity explains paradoxical findings that facial expressions can be simultaneously evaluated as both positive and negative (e.g., happily-disgusted; 10). Further, our results suggest that the use of CVML to identify individual differences in positive and negative affect recognition from dynamic facial expressions is a potentially important avenue for future research. While the current study only observed differences between three trained coders (see Figure 8), future studies may collect emotion ratings from naïve groups of participants and perform similar analyses.

Our results also provide support for the use of CVML as a valid, efficient alternative to human coders, and with further validation we expect CVML to expand the possibilities of future facial expression research in the social and behavioral sciences. For example, adoption of automatic facial coding tools will allow researchers to more easily incorporate facial expressions into models of human decision making. Decades of research show clear links between facial expressions of emotion and cognitive processes in aggregate (see 51,52), yet the dynamics between cognitive mechanisms and facial expressions are poorly understood in part due to difficulties accompanying manual coding. In fact, we are currently using computational modeling to explore cognition-expression relationships with the aid of CVML (53), which would be infeasible with manual coding of facial expressions. For example, in the current study it took less than three days to automatically extract AUs from 4,648 video recordings and train ML models to generate valence intensity ratings (using a standard desktop computer). In stark contrast, it took six months for three human coders to be trained and then code *affect intensity* across our 125 subjects—FACS coding would have taken much longer, rendering the scale of this project infeasible.

Models used in this study predicted positive emotion intensity with greater accuracy than negative emotion intensity, which may be due to the number of discrete facial actions associated with negative compared to positive emotional expressions. To support this claim, we found that importance scores for negative, but not positive, emotion ratings were spread across many different AUs and showed more variation across task instructions (Figures 5 and 6). This suggests that a wider range of facial expressions were used by coders when generating negative rather than positive emotion ratings. Future studies might address this with CVML models that can detect more than the 20 AUs used here. Additionally, our results suggest that negative affect intensity requires more training data for CVML than positive affect, as evidenced by large discrepancies in model performance between our CVML model that ignored the task instructions compared to those that we fit to data from each task instruction separately. Future studies might address this by devoting more time to collecting and coding negative, rather than positive, affective facial expressions.

Our interpretation of the computer-vision coded AUs in this study is potentially limited because we did not compare reliability of AU detection between FACET and human FACS experts. Additionally, FACET only detects 20 of the approximately 33 AUs described by FACS, so it is possible that there were other important AUs to which the human coders attended when generating valence ratings that we were unable to capture. However, our models showed excellent prediction accuracy on new data (i.e., capturing ~80% of the variance in human ratings of positive affect intensity), and we identified theoretically meaningful patterns of AUs for positive and negative emotion intensity that are consistent with prior studies (e.g., components of the *Duchenne smile*). It is unlikely that we would achieve these results if FACET did not reliably detect similar, important AUs which represented the intensity of positive and negative facial expressions produced by our 125 participants. Finally, as computer vision advances, we expect that more AUs will be easier to detect. CVML provides a scalable method that can be re-applied to previously collected facial expression recordings as technology progresses.

Although this study investigated positive and negative affect, our method could easily be extended to identify facial actions that are associated with other emotional constructs (e.g., arousal). The ability to identify specific AUs responsible for facial expression recognition has implications for various areas within the social and behavioral sciences. Opportunities may be particularly pronounced for psychopathology research, where deficits and/or biases in recognizing facial expressions of emotion are associated with a number of psychiatric disorders, including autism, alcoholism, and depression (54-56). CVML provides a framework through which both normal and abnormal emotion recognition can be studied efficiently and mechanistically, which could lead to rapid and cost-efficient markers of emotion recognition in psychopathology (57).

## Author Contributions

W.-Y. Ahn and N. Haines developed the study concept. Data were collected by M. Southward and J. Cheavens. N. Haines and W.-Y. Ahn conducted all statistical and machine learning analyses. All authors interpreted the results. N. Haines drafted the manuscript, and all authors contributed to critical manuscript revisions. All authors approved the final version of the manuscript for submission.

## Acknowledgements

We thank S. Bowman-Gibson for aiding in the manual quality check for all recordings, and J. Haaser, J. Borden, S. Choudhury, S. Okey, T. St. John, M. Stone, and S. Tolliver for manually coding videos. We also thank J. Cohn, J. Myung, A. Rogers, and H. Hahn for their comments and suggestions on previous drafts of the manuscript.

